# Increased theta/alpha synchrony in the habenula-prefrontal network with negative emotional stimuli in human patients

**DOI:** 10.1101/2020.12.30.424800

**Authors:** Yongzhi Huang, Bomin Sun, Jean Debarros, Chao Zhang, Shikun Zhan, Dianyou Li, Chencheng Zhang, Tao Wang, Peng Huang, Yijie Lai, Peter Brown, Chunyan Cao, Huiling Tan

## Abstract

Lateral habenula is believed to encode negative motivational stimuli and plays key roles in the pathophysiology of psychiatric disorders. However, how habenula activities are modulated during the perception and processing of emotional information is still poorly understood. We recorded local field potentials from bilateral habenula areas with simultaneous cortical magnetoencephalography in nine patients with psychiatric disorders during an emotional picture viewing task. Oscillatory activity in the theta/alpha band (5-10 Hz) within the habenula and prefrontal cortical regions, as well as the coupling between these structures, are increased during the perception and processing of negative emotional stimuli compared to positive emotional stimuli. The evoked increase in theta/alpha band synchronization in the frontal cortex-habenula network correlated with the emotional valence not the arousal score of the stimuli. These results provide direct evidence for increased theta/alpha synchrony within the habenula area and prefrontal cortex-habenula network in the perception of negative emotion in human participants.

## Introduction

The habenula is an epithalamic structure that functionally links the forebrain with the midbrain structures that are involved in the release of dopamine (i.e., the substantia nigra pars compacta and the ventral tegmental area) and serotonin (i.e., raphe nucleus) (Wang and Aghajanian, 1977; Herkenham and Nauta, 1979; Hikosaka *et al*., 2008; Hong *et al*., 2011; Proulx *et al*., 2014; Hu *et al*., 2020). As a region that could influence both the dopaminergic and serotonergic systems, the habenula is thought to play a key role in not only sleep and wakefulness but also in regulating various emotional and cognitive functions. Animal studies showed that activities in lateral habenula increased during the processing of aversive stimulus events such as omission of predicted rewards and stimuli provoking anxiety, stress, pain, and fear (Matsumoto and Hikosaka, 2007; Hikosaka, 2010; Yamaguchi *et al*., 2013; Hu *et al*., 2020).

Hyperexcitability and dysfunction of the lateral habenula (LHb) have been implicated in the development of psychiatric disorders including depressive disorder and bipolar disorders (Fakhoury, 2017; Yang *et al*., 2018b). In rodents, LHb firing rate and metabolism is elevated in parallel with depressive-like phenotypes such as reduction in locomotor and rearing behaviors (Caldecott-Hazard *et al*., 1988), and also increases during acquisition and recall of conditioned fear (Gonzalez-Pardo *et al*., 2012). High-resolution magnetic resonance imaging in patients has also revealed smaller habenula volume in patients with depressive and bipolar disorders (Savitz *et al*., 2011). Other evidence suggests that dysfunction of the LHb is involved in different cognitive disorders, such as schizophrenia (Shepard *et al*., 2006) and addiction (Velasquez *et al*., 2014). More direct evidence of the involvement of the LHb in psychiatric disorders in humans comes from deep brain stimulation (DBS) of the LHb that has potential therapeutic effects in treatment-resistant depression, bipolar disorder, and schizophrenia (Sartorius *et al*., 2010; Zhang *et al*., 2019; Wang *et al*., 2020). However, how habenula activities are modulated during the processing of emotional information in humans is still poorly understood.

The processing of emotional information is crucial for an individual’s mental health and has a substantial influence on social interactions and different cognitive processes. Dysfunction and dysregulation of emotion-related brain circuits may precipitate mood disorders (Phillips *et al*., 2003b). Investigating the neural activities in response to emotional stimuli in the cortical-habenula network is crucial to our understanding of emotional information processing in the brain. This might also shed light on how to modulate habenula in the treatment of psychiatric disorders. In this study, we utilize the unique opportunity offered by DBS surgery targeting habenula as a potential treatment for psychiatric disorders. We measured local field potentials (LFPs) from the habenula area using the electrodes implanted for DBS in patients during a passive emotional picture viewing task (Fig. 1; Materials and Methods). Whole brain magnetoencephalography (MEG) was simultaneously recorded. This allowed us to investigate changes in the habenula neural activity and its functional connectivity with cortical areas induced by the stimuli of different emotional valence. The high temporal resolution of the LFP and MEG measurements also allowed us to evaluate how local activities and cross-region connectivity change over time in the processing of emotional stimuli. Previous studies on rodent models of depression showed that, during the depression-like state in rodents, LHb neuron firing increased with the mean firing rate at the theta band (Li *et al*., 2011) and LHb neurons fire in bursts and phase locked to local theta band field potentials (Yang *et al*., 2018a). Therefore, we hypothesize that theta band activity in the habenula LFPs in humans would increase in response to negative emotional stimuli.

**Figure 1.**
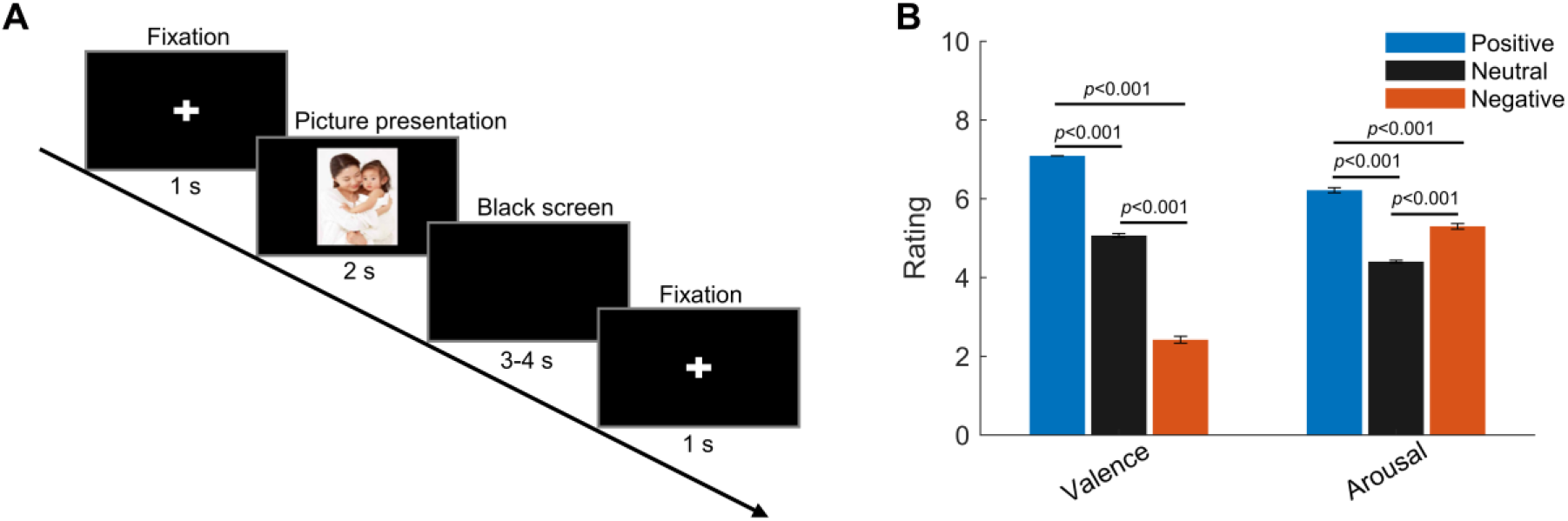
Experimental paradigm and ratings (valence and arousal) of the presented pictures. (A) Timeline of one individual trial: Each trial started with a white cross (‘+’) presented with black background for 1 second indicating the participants to get ready and pay attention; then a picture was presented in the center of the screen for 2 seconds. This was followed by a blank black screen presented for 3 to 4 second (randomized). (B) Valence and arousal ratings for figures of the three emotional categories presented to the participants. Valence: 1 = very negative; 9 = very positive; Arousal: 1 = very clam, 9 = very exciting. Error bars indicate the standard deviation of the corresponding mean across participants (N = 9).

## Results

### Spontaneous oscillatory activity in the habenula during rest includes theta/alpha activity

Electrode trajectories and contact positions of all recorded patients in this study were reconstructed using the Lead-DBS toolbox (Horn and Kuhn, 2015) and shown in Fig. 2A. The peak frequency of the oscillatory activities during rest for each electrode identified using the Fitting Oscillations and One-Over-F (FOOOF) algorithm (Haller *et al*., 2018; Donoghue *et al*., 2020) is presented in Table 1. We detected the power of oscillatory activities peaking in the theta/alpha frequency range (here defined as 5-10 Hz) in 13 out of the 18 recorded habenula during rest, compared to 7 of the 18 recorded habenula with peaks in beta band (12-30 Hz). The average peak frequency was 8.2 ± 1.1 Hz for theta/alpha, and 15.1 ± 1.8 Hz for beta band (Fig. 2B). Three out of the 18 recorded habenula showed oscillatory activities in both theta/alpha and beta bands. Fig. 2D-F shows the position of the electrodes with only theta/alpha band peaks, with only beta peaks in both sides (Case 3), with both theta/alpha and beta band peaks during rest (Case 6), respectively. The electrodes from which only alpha/theta peaks were detected are well placed in the habenula area.

**Figure 2.**
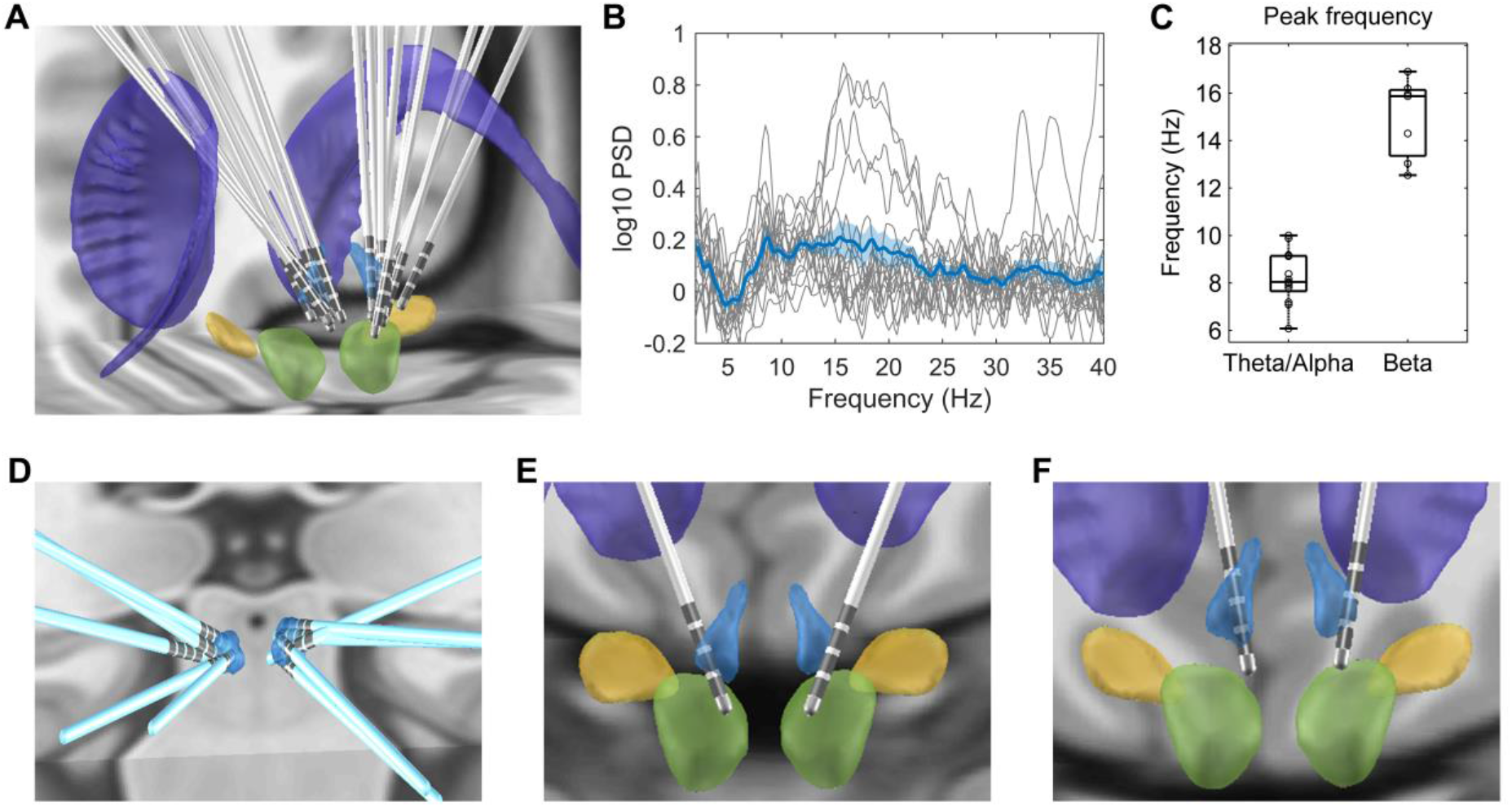
Electrode location and spectral characteristics of LFPs from recorded habenula at rest. (A) Electrode locations reconstructed using Lead-DBS, with the structures colored in light blue for the habenula, purple for the caudate nucleus, light green for the red nucleus, and yellow for subthalamic nucleus. (B) The log-transformed oscillatory power spectra fitted using *fooof* method (after removing the non-oscillatory 1/f components). The bold blue line and shadowed region indicates the mean ± SEM across all recorded hemispheres and the thin grey lines show measurements from individual hemispheres. (C) Boxplot showing the peak frequencies at theta/alpha and beta frequency bands from all recorded habenula. (D) Positions of the electrodes with theta peaks only during rest. (E) Electrode positions for Case 3, in whom only beta band peaks were detected in the resting activities from both sides. (F) Electrode positions for Case 6, in whom both theta and beta band peaks were present in resting activities from both sides.

**Table 1.** Characteristics of enrolled subjects.

| Patient | Sex | Age<br>(years) | Duration<br>(years) | Disease | HAMD<br>score | BDI<br>score | Resting oscillation peaks |  |
| --- | --- | --- | --- | --- | --- | --- | --- | --- |
|  |  |  |  |  |  |  | L | R |
| 1 | M | 21 | 5 | Schiz | NA | 32 | 9.1 Hz | 9.8 Hz |
| 2 | M | 21 | 5 | Dep | 12 | 10 | 7.9 Hz | 8.4 Hz |
| 3 | M | 44 | 10 | Bipolar | 23 | 22 | 14.3 Hz | 15.9 Hz |
| 4 | F | 19 | 4 | Schiz | NA | NA | 10 Hz | 8.1 Hz |
| 5 | M | 21 | 3 | Dep | 24 | 38 | 7.1 Hz | 16.9 Hz |
| 6 | M | 16 | 2 | Schiz | NA | 34 | 9.2 Hz; 13.0 Hz | 7.2 Hz; 12.5 Hz |
| 7 | F | 30 | 8 | Bipolar | 21 | 33 | 6.1 Hz | 7.8 Hz |
| 8 | F | 28 | 13 | Dep | 28 | 37 | No peak | 8.0 Hz |
| 9 | M | 35 | 20 | Dep | 25 | 34 | 16.2 Hz | 7.9 Hz; 16.0 Hz |

Hab, habenula; F, female; M, male; Dep, depressive disorder; Bipolar, bipolar disorder; Schiz, schizophrenia; HAMD, Hamilton Depression Rating Scale (17 items); BDI, Beck Depression Inventory. Both HAMD and BDI were acquired before the surgery. NA, not available.

### Habenular theta/alpha activity is differentially modulated by stimuli with positive and negative emotional valence

The power spectra normalized to the baseline activity (−2000 to −200 ms) showed a significant event-related synchronization (ERS) in the habenula spanning across 2-30 Hz from 50 to 800 ms after the presentation of all stimuli (*p*_cluster_ < 0.05, Fig. 3A-C). Permutation tests was applied to the power-spectra in response to the negative and positive emotional pictures from all subjects. This identified two clusters with significant difference for the two emotional valence conditions: one in the theta/alpha range (5-10 Hz) at short latency (from 100 to 500 ms, Fig. 3D and 3E) after stimulus presentation and another in the theta range (4-7 Hz) at a longer latency (from 2700 to 3300 ms, Fig. 3D and 3F), with higher increase in the identified frequency bands with negative stimuli compared to position stimuli in both clusters. The power of the activity at the identified frequency band for the neutral condition sits between the values for the negative condition and positive condition in both identified time windows (Fig. 3G-H).

**Figure 3.**
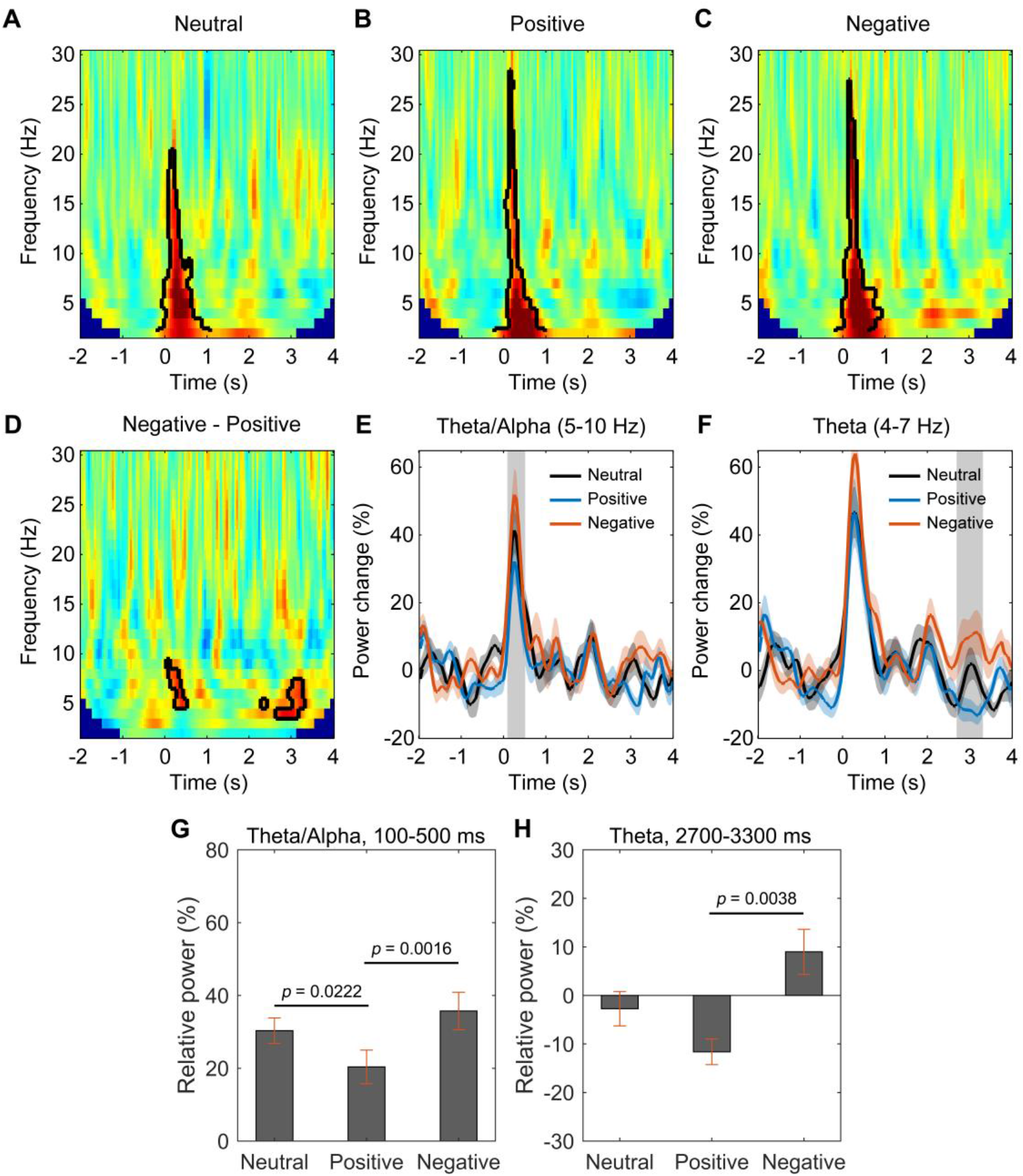
Habenular theta/alpha activity is differentially modulated by stimuli with positive and negative emotional valence (N = 18). (A-C) Time-frequency representations of the power response relative to pre-stimulus baseline (−2000 to −200 ms) for neutral (A), positive (B), and negative (C) valence stimuli, respectively. Significant clusters (*p* < 0.05, non-parametric permutation test) are encircled with a solid black line. (D) Time-frequency representation of the power response difference between negative and positive valence stimuli, showing significant increased activity the theta/alpha band (5-10 Hz) at short latency (100-500 ms) and another increased theta activity (4-7 Hz) at long latencies (2700-3300 ms) with negative stimuli (*p* < 0.05, non-parametric permutation test). (E-F) Normalized power of the activities at theta/alpha (5-10 Hz) and theta (4-7 Hz) band over time. Significant difference between the negative and positive valence stimuli is marked by a shadowed bar (*p* < 0.05, corrected for multiple comparison). (G-H) The average spectral power relative to baseline activity in the identified time period and frequency band for different emotional valence conditions (5-10 Hz, 100-500 ms; 4-7 Hz, 2700-3300 ms).

### Theta/Alpha oscillations in the prefrontal cortex are also differentially modulated by stimuli with positive and negative emotional valence

For cortical activities measured using MEG, we first computed the time-frequency power spectra normalized to the baseline activity (−2000 to −200 ms) averaged across all MEG frontal sensors highlighted in the Fig. 4A for different stimulus emotional valence conditions for each recorded participant. The average power spectra across all participants for different valence conditions are shown in Fig. 4B. Permutation test applied to the power-spectra in response to the negative and positive emotional pictures from all subjects identified clusters with significant differences (*p*_cluster_ < 0.05) in the theta/alpha range at short latency (from 100 to 500 ms after stimulus onset) (Fig. 4C). Subsequent analysis of power changes over the identified frequency band (5-10 Hz) and time window (100-500 ms) confirmed significantly increased activity with negative stimuli in frontal sensors only (Fig. 4D).

**Figure 4.**
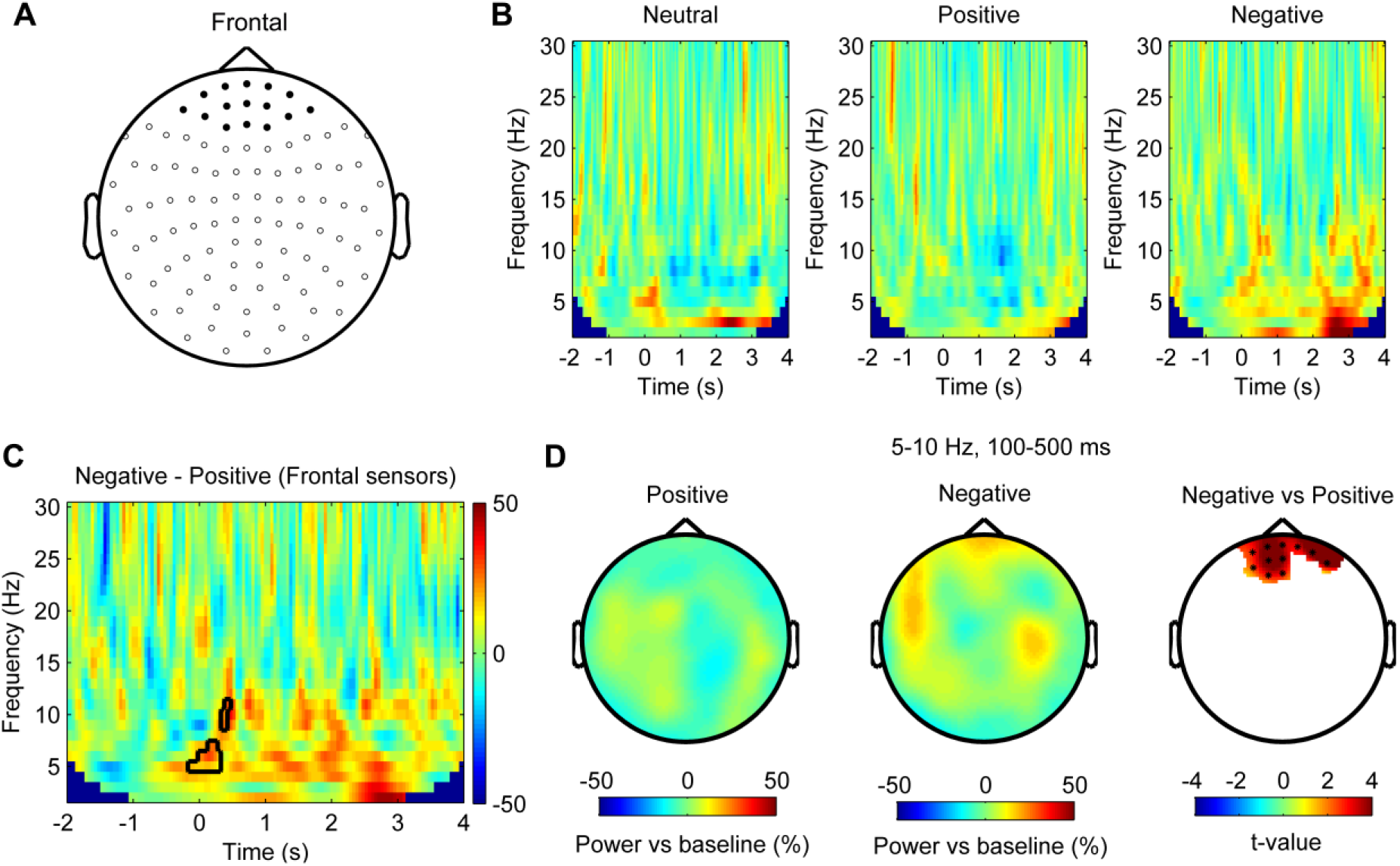
Theta/Alpha oscillations in the prefrontal cortex are differentially modulated by stimuli with positive and negative emotional valence (N = 8). (A) Layout of the MEG sensor positions and selected frontal sensors (dark spot). (B) Time-frequency representation of the power changes relative to pre-stimulus baseline for neutral, positive and negative stimuli averaged across frontal sensors (time 0 for stimuli onset). (C) Non-parametric permutation test showed clusters in the theta/alpha band at short latency after stimuli onset with significant difference (*p* < 0.05) comparing negative and positive stimuli across frontal sensors. (D) Scalp plot showing the power in the 5-10 Hz theta/alpha band activity at 100-500 ms after the onset of positive (left), negative (middle) stimuli, and statistical t-values and sensors with significant difference (right) at a 0.05 significance level (corrected for whole brain sensors).

Next, we used a frequency domain beamforming approach to identify the source of the difference in theta/alpha reactivity within the 100-500 ms time window at the corrected significance threshold of *p* < 0.05. We found two main significant source peaks with one in the right prefrontal cortex (corresponding to Brodmann area 10, MNI coordinate [16, 56, 0]; t-value = 4.14, *p* = 0.046, corrected) and the other in the left prefrontal cortex (corresponding to Brodmann area 9, MNI coordinate [-32, 38, 28]; t-value = 3.21, *p* = 0.046, corrected) (Fig. 5). No voxels within identified areas in Figure 5 showed any significant difference in the pre-cue baseline period, suggesting that the observed difference in the theta/alpha power reactivity was not due to difference in the baseline power between the two emotional valence conditions.

**Figure 5.**
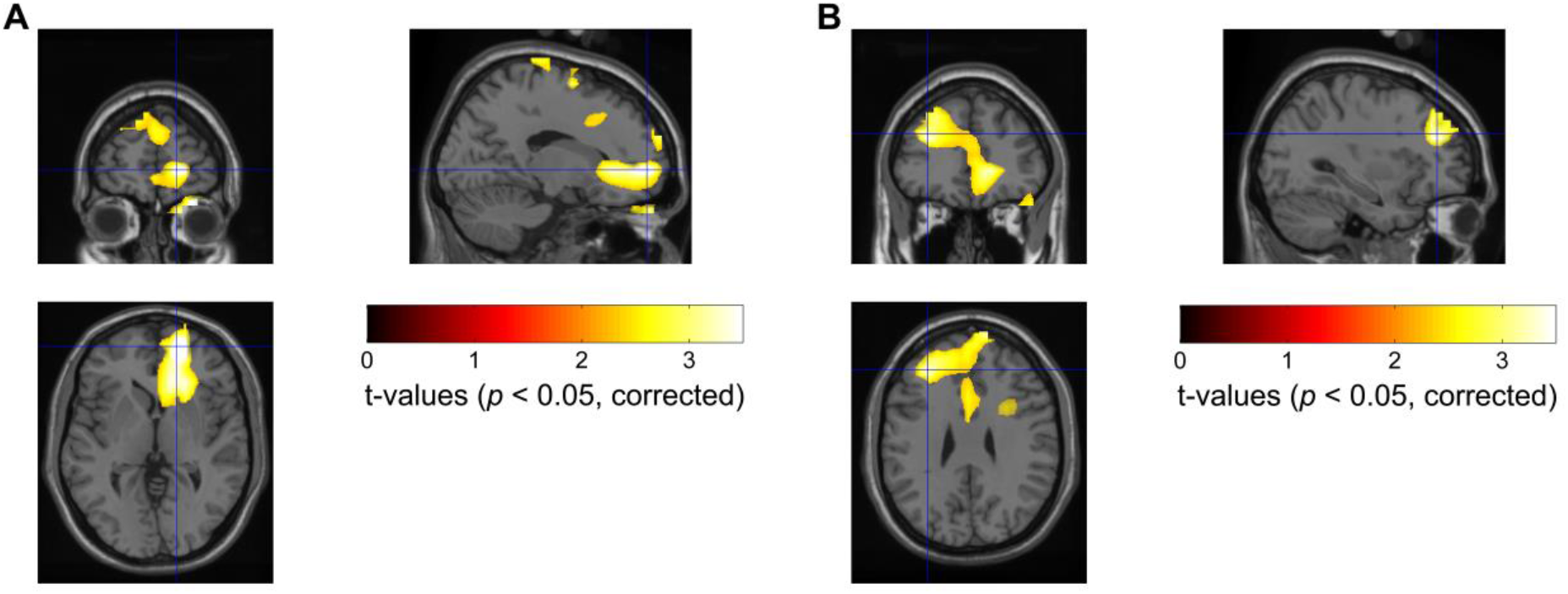
Statistical source maps of t-values (*p* < 0.05; corrected for whole brain) for the comparison of theta/alpha band (5-10 Hz) power reactivity to negative vs. positive emotional valence stimuli across subjects (N = 8). DICS beamformer was applied to the average theta/alpha band power changes from 100 to 500 ms after stimulus onset. The image was transformed to MNI template space and overlaid on the template structural image. The peak emotional valence induced differences in the theta/alpha power were localized in the right Brodmann area 10, MNI coordinate [16, 56, 0] and left Brodmann area 9, MNI coordinate [-32, 38, 28].

### Cortical-habenular coherence is also differentially modulated by stimuli with positive and negative emotional valence

In addition, we asked how the coupling between habenula and cortex in the theta/alpha activity are modulated over time in the task and how the coupling changes with the valence of the presented stimuli. The time-varying coherence between each MEG sensor and the habenula LFP was first calculated for each individual trial, and then averaged across all MEG sensors and across trials in each emotional valence condition for each habenula. Comparing the time-varying cortical-habenula coherence for the negative and positive emotional valence conditions across all recorded habenula showed increased coherence with negative stimulus in the theta/alpha band (5-10 Hz) in the time window of 800-1300 ms (N = 16, Fig. 6A). Subsequent statistical analysis of the coherence changes in this frequency band and selected time window (800-1300 ms) across the scalp revealed significant increased coherence with negative stimuli over right frontal and temporal areas (Fig. 6B). Linear mixed-effect modelling confirmed significant effect of the increase in the theta/alpha band prefrontal cortex (PFC)-habenular coherence (relative to the pre-stimulus baseline) during this time window (800-1300 ms) on the theta activity increase in the habenula at the later time window (2700-3300 ms after stimuli onset) (k = 0.2434 ± 0.1031, *p* = 0.0226, R^2^ = 0.104; Fig. 6C).

**Figure 6.**
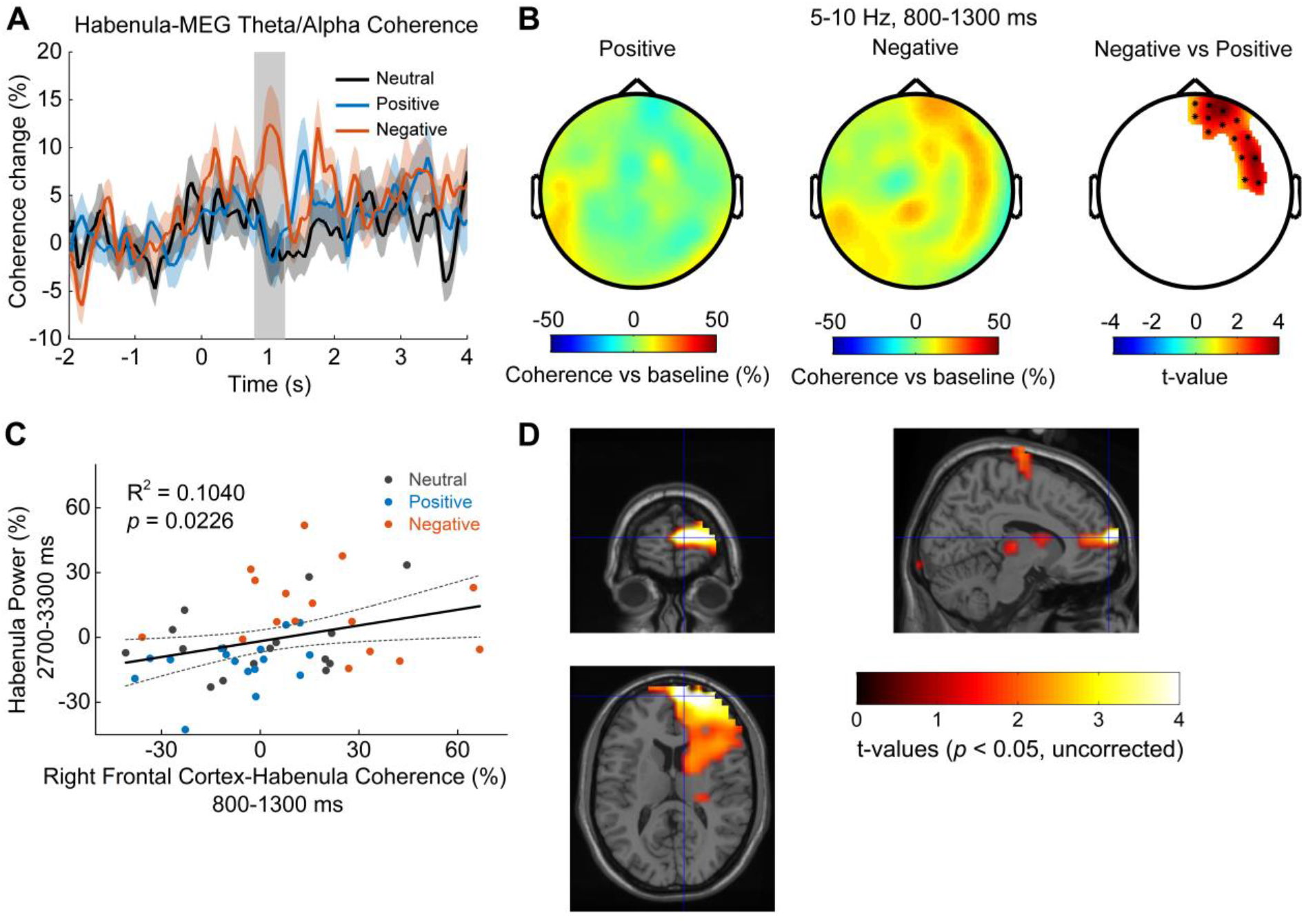
Cortical-habenular coherence in the theta/alpha band is also differentially modulated by stimuli with positive and negative emotional valence (N = 16). (A) Time-varying theta (5-10 Hz) habenula-cortical coherence changes relative to pre-cue baseline averaged across all MEG channel combinations for each recorded habenula. The thick colored lines and shaded area show the mean and standard error across all recorded habenula. The coherence was significantly higher at 800-1300 ms after the onset of negative emotional stimuli compared to positive stimuli (rectangular shadow showing the time window with *p* < 0.05). (B) Scalp plot showing the cortical-habenula coherence in the theta band during the identified time window (800-1300 ms) for positive stimuli (left), negative stimuli (middle), and statistical t-values and sensors with significant difference (right) masked at *p* < 0.05 (corrected for whole brain sensors). (C) The increase in the theta band coherence between right frontal cortex and habenula at 800-1300 ms correlated with the theta increase in habenula at 2700-3300 ms after stimuli onset; (D) Statistical source maps of t-values (*p* < 0.05; uncorrected) for the comparison of theta/alpha coherence response in the time window of 800 to 1300 ms between negative stimuli with positive stimuli. The peak coherence differences were mainly localized in the right Brodmann area 10, MNI coordinate [10, 64, 12].

Source localization of the theta/alpha habenula-cortical coherence difference for negative and positive stimuli revealed that theta/alpha coherence was higher with negative stimuli in right frontal regions, indicated in Fig. 6D. The location of the peak t-statistic (t-value = 5.73, *p* = 0.001, uncorrected) corresponds to MNI coordinate [10, 64, 12] and the region encompasses right medial prefrontal cortex.

### Increased theta/alpha synchrony in the PFC-habenula network correlated with emotional valence, not arousal

It should be noted that there was co-variation between emotional valence and arousal in the stimuli presented (Fig. 1B), and previous studies have shown that some neural activity changes in response to the viewing of affective pictures can be mediated by the effect of stimulus arousal (Huebl *et al*., 2014; Huebl *et al*., 2016). Therefore, we used linear mixed-effect modelling to assess whether the increased theta/alpha oscillations we observed in the habenula, the prefrontal cortex (PFC) and in the PFC-habenula coherence in response to the viewing of negative compared to positive emotional pictures should be attributed to the emotional valence or the stimulus arousal. The models identified significant fixed effects of valence on all the reported changes in the PFC-habenula network, but there was no effect of arousal (Table 2 for the modelling and results). The negative effects of valence indicate that the lower the emotional valence score (more negative) of the presented stimuli, the higher the theta/alpha increase within the habenula, the PFC and in the PFC-habenula theta band coherence.

**Table 2.** Linear mixed effect modelling details.

| ID | Model | Fixed Effect of<br>Valence |  | Fixed Effect of<br>Arousal |  | R<sup>2</sup> |
| --- | --- | --- | --- | --- | --- | --- |
|  |  | k-Value | p-Value | k-Value | p-Value |  |
| 1 | *HabTheta1 ~*<br>*Valence + Arousal + 1\|SubID* | -2.8044 ±<br>0.9840 | 0.0063 | -2.5221 ±<br>2.5363 | 0.3247 | 0.6191 |
| 2 | *HabTheta2 ~*<br>*Valence + Arousal + 1\|SubID* | -4.4526 ±<br>1.1753 | 0.0004 | 0.1975 ±<br>3.0295 | 0.9483 | 0.2557 |
| 3 | *PFC_Theta ~*<br>*Valence + Arousal + 1\|SubID* | -2.8921 ±<br>1.0221 | 0.0069 | -3.6237 ±<br>2.6252 | 0.1743 | 0.4368 |
| 4 | *rPFC_Hab_Coh ~*<br>*Valence + Arousal + 1\|SubID* | -6.1031 ±<br>1.6785 | 0.0007 | 3.5242 ±<br>4.3112 | 0.4180 | 0.2766 |

HabTheta1: Theta/Alpha band (5-10 Hz) in habenula LFPs at 100-500 ms
HabTheta2: Theta band (4-7 Hz) in habenula LFPs at 2700-3300 ms
PFC\_Theta: Theta/Alpha band (5-10 Hz) averaged across frontal sensors at 100-500 ms
rPFC\_Hab\_Coh: Theta/Alpha band (5-10 Hz) coherence between right PFC and habenula at 800-1300 ms
Valence: valence value for the displayed pictures (1 = unpleasant -> 5 = neutral -> 9 = pleasant)
Arousal: arousal value of the displayed pictures (1 = calm -> 9 = exciting)

## Discussion

This study has showed that oscillatory activities in the theta/alpha frequency band within the habenula and prefrontal cortical regions, as well as the connectivity between these structures in the same frequency band, are modulated in an emotional picture viewing task in human participants. Compared with positive emotional stimuli, negative emotional stimuli were associated with higher increase in theta/alpha oscillation in both habenula and bilateral frontal cortex with a short latency (from 100 to 500 ms) after stimulus onset. Furthermore, higher theta/alpha coherence between habenula and right prefrontal cortex was observed at 800 – 1300 ms after the stimulus onset, which was correlated with another increase in theta power in the habenula with a long latency (from 2700 to 3300 ms) after stimulus onset. These changes correlated with the emotional valence but not with the stimulus arousal of the presented figures. These activity changes at different time windows may reflect the different neuropsychological processes underlying emotion perception including identification and appraisal of emotional material, production of affective states, and autonomic response regulation (Phillips *et al*., 2003a). This is the first study, to our knowledge, implicating increased theta band activities in the habenula-PFC network in negative emotions in human patients.

### Habenula theta/alpha oscillations in negative emotional processing and major depression

The lateral habenula (LHb) has shown consistent hyperactivity in multiple animal models of depression-like phenotypes (Hu *et al*., 2020). Increased LHb activities have been observed during omission of a predicted reward, depressive-like phenotype, fear or stress (Matsumoto and Hikosaka, 2009; Bromberg-Martin and Hikosaka, 2011; Wang *et al*., 2017). Furthermore, manipulations enhancing or suppressing LHb activity in rodents lead to depressive-like or antidepressant effects, respectively (Li *et al*., 2013; Lecca *et al*., 2016; Cui *et al*., 2018; Yang *et al*., 2018a). Increased activation of the lateral habenula inhibits dopamine neurons (Ji and Shepard, 2007; Hikosaka, 2010) and allows avoidance of threatening or unpleasant confrontations (Shumake *et al*., 2010; Friedman *et al*., 2011). In accordance with findings in animal models, several studies have provided evidence for habenula hyperactivity in human subjects with depressive disorders (Morris *et al*., 1999; Lawson *et al*., 2017).

To our knowledge, this is the first study showing increased oscillatory activity in the habenula in the theta/alpha frequency band with perception of negative emotion in human participants. This is consistent with previous findings that LHb neurons in rodents in the depressive-like state showed increased firing with a mean firing rate in the theta frequency band (Li *et al*., 2011), and that ketamine reversed both the increase in theta activity in the habenula and depressive-like behavior in rodents (Yang *et al*., 2018a). The results in this study are also consistent with recent research showing that acute 5 Hz deep brain stimulation of the lateral habenula is associated with depressive-like behavior such as increased duration of immobility in a forced swim test in rodents (Jakobs *et al*., 2019). Possibly due to the limited sample size, we didn’t observe any correlation between the habenula theta/alpha activities and the Beck Depression Inventory score or Hamilton Depression Rating Scale score measured before the surgery across patients in this study. It therefore remains to be established whether hyper-synchrony in the theta band in habenula might be associated with the development of depressive symptoms in human patients.

### Prefrontal cortex-habenular coherence in negative emotional processing

Apart from increased theta/alpha band synchronization within the bilateral habenula and prefrontal cortex, our data showed that negative emotional stimuli induced increased theta/alpha coherence between the habenula and the right prefrontal cortex. The increased rPFC-habenular coherence correlated with further increase of theta activities within the habenula at a later latency. These results suggest a specific role of the theta/alpha synchronization between habenula and frontal cortex in the perception of negative emotional valence. Previous studies have showed that LHb receives input from cortical areas processing information about pain, loss, adversities, bad, harmful or suboptimal choices, such as the anterior insula and dorsal ACC (dACC) and the pregenual ACC (pgACC) (Vadovicova, 2014). Our data is consistent with the hypothesis that PFC-to-habenular projections provide a teaching signal for value-based choice behavior, helping to learn to avoid potentially harmful, low valued or wrong choices (Vadovicova, 2014).

Our data also showed that the increase of PFC-habenular coherence during the presentation of negative emotional stimuli were mainly located in the right frontal cortex. Many studies have investigated how both hemispheres have a role in emotional processing. The Right Hemisphere Hypothesis (RHH) suggests that the right hemisphere would be involved, more than the left hemisphere, in the processing of all emotional stimuli, irrespective of their emotional valence (Gainotti, 2012). On the other hand, the Valence Hypothesis (VH) posits that the left and the right hemispheres would be specialized in processing positive and negative emotions, respectively (Davidson, 1992; Wyczesany *et al*., 2018). The latter hypothesis has also been supported by studies of brain lesion (Starkstein *et al*., 1987; Morris *et al*., 1999), electroencephalography (EEG) (Davidson, 1992; Wyczesany *et al*., 2018), transcranial magnetic stimulation (TMS) (Pascual-Leone *et al*., 1996) and functional neuroimaging (Canli *et al*., 1998; Beraha *et al*., 2012) in the prefrontal cortex. Our findings suggest a more important role of the functional connectivity between the right frontal cortex and habenula for the processing of negative emotions.

### Implications for the development of DBS therapy

Although the exact underlying physiological mechanism of DBS remains elusive, high frequency DBS delivered to STN and GPi can reduce the firing rates of local neurons (Boraud *et al*., 1996; Welter *et al*., 2004) and suppress the hypersynchrony of oscillatory activities in the beta frequency band in the network leading to symptom alleviation (Kuhn *et al*., 2008; Oswal *et al*., 2016) for Parkinson’s disease. In addition, high frequency DBS may also dissociate input and output signals, resulting in the disruption of abnormal information flow through the stimulation site (Chiken and Nambu, 2016). This is supported by recent studies showing that patient specific connectivity profiles between the stimulation target and area of interest in the cortex can predict clinical outcome of deep brain stimulation for Parkinson disease (Horn *et al*., 2017), major depressive disorder (MDD) (Riva-Posse *et al*., 2014) and obsessive-compulsive disorder (Baldermann *et al*., 2019). Our results suggest that increased theta oscillatory activity in the habenula and increased theta/alpha coherence between prefrontal cortex and habenula are associated with negative emotional valence in human patients. High frequency DBS targeting habenula may be beneficial for treatment-resistant MDD by inhibiting possible hyperactivity and theta band over-synchrony of neuronal activities in the habenula, and by disrupting the information flow from the prefrontal cortex to other midbrain areas through the habenula. It remains to be explored whether theta band synchronization can be used as a biomarker for closed loop habenula deep brain stimulation for better treatment of MDD.

### Limitations

The response to emotional tasks is likely to be altered in patients with pathological mood states compared to healthy subjects. This study cannot address whether the emotional valence effect we observed is specific to psychiatric disorders or is a common feature of healthy emotional processing. Another caveat we would like to acknowledge is that the human habenula is a small region compared to the size of the electrode contact used for recording (Fig. 2A). Considering that the location of habenula is adjacent to the posterior end of the medial dorsal thalamus, we may have captured activities from the medial dorsal thalamus. In addition, it should also be noted that a post-operative stun effect cannot be excluded, which could interfere with neural recordings, considering that the experiment took place only a few days after electrode implantation.

### Conclusion

In this study, we exploited the high temporal resolution of LFP and MEG measurements and observed an emotional valence effect in local activities and in cross-region coherence in the cortical-habenula network in different time windows. Our results provide evidence for the role of oscillatory activity in the theta/alpha frequency band within the habenula and prefrontal cortical regions, as well as of theta/alpha coherence between these structures in the processing and experiencing of negative emotions in human patients.

## Materials and Methods

### Participants

Nine patients (6 males, aged 16 – 44, more details in Table 1) were recruited for this study, who underwent bilateral DBS surgery targeting the habenula as a clinical trial for treatment-resistant major depression (ClinicalTrials.gov Identifier: NCT03347487) or as a pilot study for intractable schizophrenia or bipolar disorders. All participants gave written informed consent to the current study, which was approved by the local ethics committee of Ruijin hospital, Shanghai Jiao Tong University School of Medicine in accordance with the declaration of Helsinki. The surgical procedure has been previously described (Zhang *et al*., 2019). The electrode position, stimulation parameters and clinical outcome in Case 1 have been separately reported (Wang *et al*., 2020).

### Deep Brain Stimulation Operation

Implantation of the quadripolar DBS electrodes (model 3389 (contact: 1.5 mm, distance: 0.5mm, diameter: 1.27 mm); Medtronic, Minneapolis, MN, USA) was performed under general anesthesia bilaterally using a MRI-guided targeting (3.0 T, General Electric, Waukesha, WI, USA). The MRI was co-registered with a CT image (General Electric, Waukesha, WI, USA) with the Leksell stereotactic frame to obtain the coordinate values (Zhang *et al*., 2019). The electrode leads were temporary externalized for one week.

### Paradigm

Patients were recorded in an emotional picture viewing task (Kuhn *et al*., 2005; Huebl *et al*., 2016) 2 – 5 days after the first stage of the surgery for electrode implantation and prior to the second operation to connect the electrode to the subcutaneous pulse generator. During the task, participants were seated in the MEG scanner with a displaying monitor in front of them. Pictures selected from the Chinese Affective Pictures System (CAPS) (Bai *et al*., 2005) were presented on the monitor in front of them. The emotional valence (1=unpleasant ⇒ 5=neutral ⇒ 9=pleasant) and arousal (1 = calm 9 = exciting) of the pictures were previously rated by healthy Chinese participants (Bai *et al*., 2005). The figures can be classified into three valence categories (neutral, positive and negative) according to the average score on emotional valence. In our paradigm, each experiment consisted of multiple blocks of 30 trials, with each block including 10 pictures of each valence category (neutral, positive and negative) in randomized order. Each trial started with a white cross (‘+’) presented with a black background for 1 second indicating the participants to get ready and pay attention, then a picture was presented in the center of the screen for 2 seconds. This was followed by a blank black screen presented for 3 to 4 second (randomized). The task was programed using PsychoPy (https://www.psychopy.org/) with the timeline of each individual trial shown in Fig. 1A. The participants were reminded to pay attention to the pictures displayed on the monitor and they were instructed to try to experience the emotions the pictures conveyed. An additional neutral picture was presented randomly three times per block, upon which the patients were supposed to press a button to ensure constant attention during the paradigm. All participants completed 2 to 4 blocks of the paradigm and none of them missed any response to the additional figure indicating that they kept focus and that their working memory required for the task is normal. Pictures displayed to different participants are overlapped but not exactly the same, the average valence and arousal values of the displayed pictures are as shown in Fig. 1B. There were significant differences in the emotional valence scores, as well as in the arousal scores for the presented figures of the three emotional valence categories (one-way ANOVA followed by Bonferroni post hoc test, F_2,24_ = 14642.02, *p* < 0.0001 for the valence score, and F_2,24_ = 2102.55, *p* < 0.0001 for the arousal score). The positive figures have the highest valence scores and highest arousal scores; the negative figures have the lowest valence scores; whereas the neutral figures have lowest arousal scores.

### Data Acquisition

Whole-brain MEG and LFP were simultaneously recorded at a sampling frequency of 1000 Hz using a 306-channel, whole-head MEG system with integrated EEG channels (Elekta Oy, Helsinki, Finland). LFPs from all individual contacts (0, 1, 2, and 3, with 0 being the deepest contact) of the DBS electrodes were measured in monopolar mode with reference to a surface electrode attached to the earlobe or one of the most dorsal DBS contact. The MaxFilter software (Elekta Oy, Helsinki, Finland) was used to apply the temporally extended signal space separation method (tSSS) to the original MEG data for removing the magnetic artefacts and movement artefacts (Taulu and Simola, 2006). The MEG and LFP recordings were synchronized with the timing of the onset of each picture stimuli through an analogue signal sent by the laptop running the picture viewing paradigm. The voltage of the analogue signal increased at the onset of the presentation of each picture and lasted for 500 ms before going back to zero. The voltage increase was different for pictures of different emotional valence category.

### Reconstruction of Electrode Locations in the Habenula

We used the Lead-DBS toolbox (Horn and Kuhn, 2015) to reconstruct the electrode trajectories and contact locations for all recorded patients (Fig. 2A). Post-operative CT was co-registered to pre-operative T1 MRI using a two-stage linear registration as implemented in Advanced Normalization Tools (ANT) (Avants *et al*., 2008). CT and MRI were spatially normalized into MNI_ICBM_2009b_NLIN_ASYM space (Fonov *et al*., 2011). Electrodes were automatically pre-localized in native and template space using the PaCER algorithm (Husch *et al*., 2018) and then manually localized based on post-operative CT (Horn and Kuhn, 2015).

### LFP and MEG Data Analysis

All data were analyzed using Matlab (R2013b) with FieldTrip (version 20170628) (Oostenveld *et al*., 2011) and SPM8 toolboxes. Bipolar LFP recordings were constructed offline by subtracting the monopolar recordings from neighboring contacts on each electrode. One bipolar LFP channel within or closest to the habenula was selected from each recorded hemisphere for final analysis based on the post-operative imaging data and the location reconstruction based on Lead-DBS (Fig. 2A). Artefacts due to movement, flat and jump artefacts were visually inspected and manually marked during the pre-processing with FieldTrip. All the selected bipolar LFPs and MEG recordings were high-pass filtered at 0.3 Hz, notch-filtered at 50 Hz and higher-order harmonics, low pass filtered at 100 Hz and then down-sampled to 250 Hz before further analysis. Eye blink and heartbeat artefacts in the MEG signals were identified by ICA and the low frequency, high amplitude components were removed from all MEG sensors. The MEG data of one subject (case 4) had to be discarded due to severe artefacts across all MEG channels. Hence, all reported results with MEG data are based on eight subjects.

The oscillatory activities in the habenula LFPs during rest were first investigated. The power spectra were calculated using the Fast Fourier Transform (FFT). We then applied the Fitting Oscillations and One-Over-F (FOOOF) algorithm (Haller *et al*., 2018; Donoghue *et al*., 2020) to separate the LFP power spectral densities into aperiodic (1/f-like component) and periodic oscillatory components which are modelled as Gaussian peaks. With this algorithm, a periodic oscillatory component is detected only when its peak power exceeds that of aperiodic activity by a specified threshold. In this study, the algorithm was applied to the 2-40 Hz range of the raw power spectra of the LFPs from each recorded hemisphere. We set the maximal number of power peaks (max_n_peaks) to be four, the width of the oscillatory peak (peak_width_limits) to be between 1 and 15, and the threshold for detecting the peak (peak_threshold) to be 2. The goodness of fit was visually inspected for recordings from each hemisphere to make sure that the parameter settings worked well. After removing the aperiodic component, the periodic oscillatory components in the LFP power spectra were parameterized by their center frequency (defined as the mean of the Gaussian), amplitude (defined as the distance between the peak of the Gaussian and the aperiodic fit), and bandwidth (defined as two standard deviations of the fitted Gaussian) of the power peaks.

In the next step, we investigated the event-related power changes in the habenula LFPs and MEG signals in response to the presentation of figures of different emotional valence categories. All LFP and MEG signals were divided into event-related epochs aligned to the stimuli onset (−2500 to 4500 ms around the stimulus onset) and visually inspected for artefacts due to movement and other interferences. Trials with artefacts were removed from final analysis, leaving a mean number of 27 trials (range 18 – 30) for each valence category for each subject. A time-frequency decomposition using the wavelet transform-based approach with Morlet wavelet and cycle number of 6 was applied to each trial. We used a 500 ms buffer on both sides of the clipped data to reduce edge effects. The time-frequency representations were then averaged across trials of the same valence condition and baseline corrected to the average of pre-stimulus activity (−2000 to −200 ms) for each frequency band. Thus, resulting time-frequency values were percentage changes in power relative to the pre-stimulus baseline.

### MEG-specific Data Analysis

Statistical comparison of power over a determined frequency band and time window between stimulus conditions across the group of subjects was performed to find topographical space difference. MEG source localization was conducted using a frequency domain beamforming approach. The dynamic imaging of coherent sources (DICS) beamformer in SPM8 with a single-shell forward model was used to generate maps of the source power difference between conditions on a 5 mm grid co-registered to MNI coordinates (Gross *et al*., 2001). In this study, we focused our source analysis on the frequency band and time window identified by previous sensor-level power analysis to locate cortical sources of significant difference in the power response to negative and positive emotional stimuli.

### Cortical-Habenular Connectivity

The functional connectivity between habenula and cortical areas was investigated using coherence analysis, which provides a frequency domain measure of the degree of co-variability between signals (Litvak *et al*., 2010; Neumann *et al*., 2015). First, time-resolved coherence in the theta/alpha frequency band between the habenula LFP and each MEG channel at sensor level were calculated using the wavelet transform-based approach with Morlet wavelet and cycle number of 6. Secondly, we determined the time window of interest by statistically comparing the sensor-level coherence between stimulus conditions. Third, cortical sources coherent with habenula-LFP activity in the determined frequency band and time window were located using DICS beamformer for each stimuli condition (Gross *et al*., 2001; Litvak *et al*., 2011).

### Statistics

A non-parametric cluster-based permutation approach (Maris and Oostenveld, 2007) was applied to normalized time-frequency matrices to identify clusters (time window and frequency band) with significant differences in the power changes induced by the presentation of pictures of different emotional valence. To achieve this, the original paired samples were randomly permuted 1000 times such that each pair was maintained but its assignment to the condition (negative or positive) may have changed to create a null-hypothesis distribution. For each permutation, the sum of the z-scores within suprathreshold-clusters (pre-cluster threshold: *p* < 0.05) was computed to obtain a distribution of the 1000 largest suprathreshold-cluster values. If the sum of the z-scores within a suprathreshold-cluster of the original difference exceeded the 95th percentile of the permutation distribution, it was considered statistically significant. The average powers in the determined frequency band and time window identified by the cluster-based permutation method between different valence conditions were further compared using post-hoc paired t-test. A one-tailed dependent-sample t statistics and cluster-based permutation testing was applied to statistically quantify the differences in DICS source for power or source coherence between negative and positive emotional stimuli. In addition, linear mixed-effect modelling (‘fitlme’ in Matlab) with different recorded subjects as random effects was used to investigate the correlations between the observed changes in the neural signals and to investigate whether any changes we observed in the neural activities were related to the ratings of the emotional valence or stimulus arousal of the stimuli. The estimated mean value and standard error of the fixed effect and associated p values, as well as the R^2^ value of the model were reported.

## Acknowledgements

We would like thank Dr Wolf-Julian Neumann at Charité–University Medicine Berlin for the discussion on the paradigm and Yingying Zhang in the help of the emotion scaling.

## References

Avants BB, Epstein CL, Grossman M, Gee JC. Symmetric diffeomorphic image registration with cross-correlation: evaluating automated labeling of elderly and neurodegenerative brain. Med Image Anal 2008; 12(1): 26–41.

Bai L, Ma H, Huang Y-X. The development of native Chinese affective picture system-a pretest in 46 college students. Chinese Mental Health J 2005; 11: 719–22.

Baldermann JC, Melzer C, Zapf A, Kohl S, Timmermann L, Tittgemeyer M, et al. Connectivity Profile Predictive of Effective Deep Brain Stimulation in Obsessive-Compulsive Disorder. Biol Psychiatry 2019; 85(9): 735–43.

Beraha E, Eggers J, Hindi Attar C, Gutwinski S, Schlagenhauf F, Stoy M, et al. Hemispheric asymmetry for affective stimulus processing in healthy subjects--a fMRI study. PloS one 2012; 7(10): e46931.

Boraud T, Bezard E, Bioulac B, Gross C. High frequency stimulation of the internal Globus Pallidus (GPi) simultaneously improves parkinsonian symptoms and reduces the firing frequency of GPi neurons in the MPTP-treated monkey. Neurosci Lett 1996; 215(1): 17–20.

Bromberg-Martin ES, Hikosaka O. Lateral habenula neurons signal errors in the prediction of reward information. Nat Neurosci 2011; 14(9): 1209–16.

Caldecott-Hazard S, Mazziotta J, Phelps M. Cerebral correlates of depressed behavior in rats, visualized using 14C-2-deoxyglucose autoradiography. J Neurosci 1988; 8(6): 1951–61.

Canli T, Desmond JE, Zhao Z, Glover G, Gabrieli JD. Hemispheric asymmetry for emotional stimuli detected with fMRI. Neuroreport 1998; 9(14): 3233–9.

Chiken S, Nambu A. Mechanism of Deep Brain Stimulation: Inhibition, Excitation, or Disruption? Neuroscientist 2016; 22(3): 313–22.

Cui Y, Yang Y, Ni Z, Dong Y, Cai G, Foncelle A, et al. Astroglial Kir4.1 in the lateral habenula drives neuronal bursts in depression. Nature 2018; 554(7692): 323–7.

Davidson RJ. Anterior cerebral asymmetry and the nature of emotion. Brain Cogn 1992; 20(1): 125–51.

Donoghue T, Haller M, Peterson EJ, Varma P, Sebastian P, Gao R, et al. Parameterizing neural power spectra into periodic and aperiodic components. Nat Neurosci 2020; 23(12): 1655–65.

Fakhoury M. The habenula in psychiatric disorders: More than three decades of translational investigation. Neurosci Biobehav Rev 2017; 83: 721–35.

Fonov V, Evans AC, Botteron K, Almli CR, McKinstry RC, Collins DL, et al. Unbiased average age-appropriate atlases for pediatric studies. NeuroImage 2011; 54(1): 313–27.

Friedman A, Lax E, Dikshtein Y, Abraham L, Flaumenhaft Y, Sudai E, et al. Electrical stimulation of the lateral habenula produces an inhibitory effect on sucrose self-administration. Neuropharmacology 2011; 60(2-3): 381–7.

Gainotti G. Unconscious processing of emotions and the right hemisphere. Neuropsychologia 2012; 50(2): 205–18.

Gonzalez-Pardo H, Conejo NM, Lana G, Arias JL. Different brain networks underlying the acquisition and expression of contextual fear conditioning: a metabolic mapping study. Neuroscience 2012; 202: 234–42.

Gross J, Kujala J, Hamalainen M, Timmermann L, Schnitzler A, Salmelin R. Dynamic imaging of coherent sources: Studying neural interactions in the human brain. Proc Natl Acad Sci U S A 2001; 98(2): 694–9.

Haller M, Donoghue T, Peterson E, Varma P, Sebastian P, Gao R, et al. Parameterizing neural power spectra. BioRxiv 2018: 299859.

Herkenham M, Nauta WJ. Efferent connections of the habenular nuclei in the rat. J Comp Neurol 1979; 187(1): 19–47.

Hikosaka O. The habenula: from stress evasion to value-based decision-making. Nat Rev Neurosci 2010; 11(7): 503–13.

Hikosaka O, Sesack SR, Lecourtier L, Shepard PD. Habenula: crossroad between the basal ganglia and the limbic system. J Neurosci 2008; 28(46): 11825–9.

Hong S, Jhou TC, Smith M, Saleem KS, Hikosaka O. Negative reward signals from the lateral habenula to dopamine neurons are mediated by rostromedial tegmental nucleus in primates. J Neurosci 2011; 31(32): 11457–71.

Horn A, Kuhn AA. Lead-DBS: a toolbox for deep brain stimulation electrode localizations and visualizations. NeuroImage 2015; 107: 127–35.

Horn A, Reich M, Vorwerk J, Li N, Wenzel G, Fang Q, et al. Connectivity Predicts deep brain stimulation outcome in Parkinson disease. Ann Neurol 2017; 82(1): 67–78.

Hu H, Cui Y, Yang Y. Circuits and functions of the lateral habenula in health and in disease. Nat Rev Neurosci 2020; 21(5): 277–95.

Huebl J, Brucke C, Merkl A, Bajbouj M, Schneider GH, Kuhn AA. Processing of emotional stimuli is reflected by modulations of beta band activity in the subgenual anterior cingulate cortex in patients with treatment resistant depression. Soc Cogn Affect Neurosci 2016; 11(8): 1290–8.

Huebl J, Spitzer B, Brucke C, Schonecker T, Kupsch A, Alesch F, et al. Oscillatory subthalamic nucleus activity is modulated by dopamine during emotional processing in Parkinson’s disease. Cortex 2014; 60: 69–81.

Husch A, M VP, Gemmar P, Goncalves J, Hertel F. PaCER - A fully automated method for electrode trajectory and contact reconstruction in deep brain stimulation. Neuroimage Clin 2018; 17: 80–9.

Jakobs M, Pitzer C, Sartorius A, Unterberg A, Kiening K. Acute 5Hz deep brain stimulation of the lateral habenula is associated with depressive-like behavior in male wild-type Wistar rats. Brain Res 2019; 1721: 146283.

Ji H, Shepard PD. Lateral habenula stimulation inhibits rat midbrain dopamine neurons through a GABA(A) receptor-mediated mechanism. J Neurosci 2007; 27(26): 6923–30.

Kuhn AA, Hariz MI, Silberstein P, Tisch S, Kupsch A, Schneider GH, et al. Activation of the subthalamic region during emotional processing in Parkinson disease. Neurology 2005; 65(5): 707–13.

Kuhn AA, Kempf F, Brucke C, Gaynor Doyle L, Martinez-Torres I, Pogosyan A, et al. High-frequency stimulation of the subthalamic nucleus suppresses oscillatory beta activity in patients with Parkinson’s disease in parallel with improvement in motor performance. J Neurosci 2008; 28(24): 6165–73.

Lawson RP, Nord CL, Seymour B, Thomas DL, Dayan P, Pilling S, et al. Disrupted habenula function in major depression. Mol Psychiatry 2017; 22(2): 202–8.

Lecca S, Pelosi A, Tchenio A, Moutkine I, Lujan R, Herve D, et al. Rescue of GABAB and GIRK function in the lateral habenula by protein phosphatase 2A inhibition ameliorates depression-like phenotypes in mice. Nat Med 2016; 22(3): 254–61.

Li B, Piriz J, Mirrione M, Chung C, Proulx CD, Schulz D, et al. Synaptic potentiation onto habenula neurons in the learned helplessness model of depression. Nature 2011; 470(7335): 535–9.

Li K, Zhou T, Liao L, Yang Z, Wong C, Henn F, et al. betaCaMKII in lateral habenula mediates core symptoms of depression. Science 2013; 341(6149): 1016–20.

Litvak V, Eusebio A, Jha A, Oostenveld R, Barnes GR, Penny WD, et al. Optimized beamforming for simultaneous MEG and intracranial local field potential recordings in deep brain stimulation patients. NeuroImage 2010; 50(4): 1578–88.

Litvak V, Jha A, Eusebio A, Oostenveld R, Foltynie T, Limousin P, et al. Resting oscillatory cortico-subthalamic connectivity in patients with Parkinson’s disease. Brain 2011; 134(Pt 2): 359–74.

Maris E, Oostenveld R. Nonparametric statistical testing of EEG- and MEG-data. J Neurosci Methods 2007; 164(1): 177–90.

Matsumoto M, Hikosaka O. Lateral habenula as a source of negative reward signals in dopamine neurons. Nature 2007; 447(7148): 1111–5.

Matsumoto M, Hikosaka O. Representation of negative motivational value in the primate lateral habenula. Nat Neurosci 2009; 12(1): 77–84.

Morris JS, Smith KA, Cowen PJ, Friston KJ, Dolan RJ. Covariation of activity in habenula and dorsal raphe nuclei following tryptophan depletion. NeuroImage 1999; 10(2): 163–72.

Neumann WJ, Jha A, Bock A, Huebl J, Horn A, Schneider GH, et al. Cortico-pallidal oscillatory connectivity in patients with dystonia. Brain 2015; 138(Pt 7): 1894–906.

Oostenveld R, Fries P, Maris E, Schoffelen JM. FieldTrip: Open source software for advanced analysis of MEG, EEG, and invasive electrophysiological data. Comput Intell Neurosci 2011; 2011: 156869.

Oswal A, Beudel M, Zrinzo L, Limousin P, Hariz M, Foltynie T, et al. Deep brain stimulation modulates synchrony within spatially and spectrally distinct resting state networks in Parkinson’s disease. Brain 2016; 139(Pt 5): 1482–96.

Pascual-Leone A, Catala MD, Pascual-Leone Pascual A. Lateralized effect of rapid-rate transcranial magnetic stimulation of the prefrontal cortex on mood. Neurology 1996; 46(2): 499–502.

Phillips ML, Drevets WC, Rauch SL, Lane R. Neurobiology of emotion perception I: The neural basis of normal emotion perception. Biol Psychiatry 2003a; 54(5): 504–14.

Phillips ML, Drevets WC, Rauch SL, Lane R. Neurobiology of emotion perception II: Implications for major psychiatric disorders. Biol Psychiatry 2003b; 54(5): 515–28.

Proulx CD, Hikosaka O, Malinow R. Reward processing by the lateral habenula in normal and depressive behaviors. Nat Neurosci 2014; 17(9): 1146–52.

Riva-Posse P, Choi KS, Holtzheimer PE, McIntyre CC, Gross RE, Chaturvedi A, et al. Defining critical white matter pathways mediating successful subcallosal cingulate deep brain stimulation for treatment-resistant depression. Biol Psychiatry 2014; 76(12): 963–9.

Sartorius A, Kiening KL, Kirsch P, von Gall CC, Haberkorn U, Unterberg AW, et al. Remission of major depression under deep brain stimulation of the lateral habenula in a therapy-refractory patient. Biol Psychiatry 2010; 67(2): e9–e11.

Savitz JB, Nugent AC, Bogers W, Roiser JP, Bain EE, Neumeister A, et al. Habenula volume in bipolar disorder and major depressive disorder: a high-resolution magnetic resonance imaging study. Biol Psychiatry 2011; 69(4): 336–43.

Shepard PD, Holcomb HH, Gold JM. Schizophrenia in translation: the presence of absence: habenular regulation of dopamine neurons and the encoding of negative outcomes. Schizophr Bull 2006; 32(3): 417–21.

Shumake J, Ilango A, Scheich H, Wetzel W, Ohl FW. Differential neuromodulation of acquisition and retrieval of avoidance learning by the lateral habenula and ventral tegmental area. J Neurosci 2010; 30(17): 5876–83.

Starkstein SE, Robinson RG, Price TR. Comparison of cortical and subcortical lesions in the production of poststroke mood disorders. Brain 1987; 110 (Pt 4): 1045–59.

Taulu S, Simola J. Spatiotemporal signal space separation method for rejecting nearby interference in MEG measurements. Phys Med Biol 2006; 51(7): 1759–68.

Vadovicova K. Affective and cognitive prefrontal cortex projections to the lateral habenula in humans. Front Hum Neurosci 2014; 8: 819.

Velasquez KM, Molfese DL, Salas R. The role of the habenula in drug addiction. Front Hum Neurosci 2014; 8: 174.

Wang D, Li Y, Feng Q, Guo Q, Zhou J, Luo M. Learning shapes the aversion and reward responses of lateral habenula neurons. eLife 2017; 6.

Wang RY, Aghajanian GK. Physiological evidence for habenula as major link between forebrain and midbrain raphe. Science 1977; 197(4298): 89–91.

Wang Y, Zhang C, Zhang Y, Gong H, Li J, Jin H, et al. Habenula deep brain stimulation for intractable schizophrenia: a pilot study. Neurosurg Focus 2020; 49(1): E9.

Welter ML, Houeto JL, Bonnet AM, Bejjani PB, Mesnage V, Dormont D, et al. Effects of high-frequency stimulation on subthalamic neuronal activity in parkinsonian patients. Arch Neurol 2004; 61(1): 89–96.

Wyczesany M, Capotosto P, Zappasodi F, Prete G. Hemispheric asymmetries and emotions: Evidence from effective connectivity. Neuropsychologia 2018; 121: 98–105.

Yamaguchi T, Danjo T, Pastan I, Hikida T, Nakanishi S. Distinct roles of segregated transmission of the septo-habenular pathway in anxiety and fear. Neuron 2013; 78(3): 537–44.

Yang Y, Cui Y, Sang K, Dong Y, Ni Z, Ma S, et al. Ketamine blocks bursting in the lateral habenula to rapidly relieve depression. Nature 2018a; 554(7692): 317–22.

Yang Y, Wang H, Hu J, Hu H. Lateral habenula in the pathophysiology of depression. Curr Opin Neurobiol 2018b; 48: 90–6.

Zhang C, Kim SG, Li D, Zhang Y, Li Y, Husch A, et al. Habenula deep brain stimulation for refractory bipolar disorder. Brain Stimul 2019; 12(5): 1298–300.

